# Rigidity, normal modes and flexible motion of a SARS-CoV-2 (COVID-19) protease structure

**DOI:** 10.1101/2020.03.10.986190

**Authors:** Stephen A. Wells

## Abstract

The rigidity and flexibility of two recently reported crystal structures (PDB entries 6Y2E and 6LU7) of a protease from the SARS-CoV-2 virus, the infectious agent of the COVID-19 respiratory disease, has been investigated using pebble-game rigidity analysis, elastic network model normal mode analysis, and all-atom geometric simulations. This computational investigation of the viral protease follows protocols that have been effective in studying other homodimeric enzymes. The protease is predicted to display flexible motions in vivo which directly affect the geometry of a known inhibitor binding site and which open new potential binding sites elsewhere in the structure. A database of generated PDB files representing natural flexible variations on the crystal structures has been produced and made available for download from an institutional data archive. This information may inform structure-based drug design and fragment screening efforts aimed at identifying specific antiviral therapies for the treatment of COVID-19.

## Introduction

As of late 2019, there has been global concern over a novel respiratory disease (designated COVID-19 by the World Health Organisation (WHO)) which originated in the Wuhan region of China and has subsequently spread to multiple other countries worldwide. The virus causing COVID-19 is a coronavirus with the taxonomic identifier SARS-CoV-2[1]. Thanks to the diligent efforts of structural biologists, several crystal structures of proteins from SARS-CoV-2 have already been obtained and made available through the Protein Data Bank (PDB) even in advance of journal publication.

Crystal structures provide us with detailed structural knowledge of the arrangement of atoms making up a protein or protein complex. A crystal structure is a static snapshot of the protein “frozen” into one of its many possible conformations. This static snapshot is rich in implicit information on the natural dynamics which the protein will explore *in vivo*. These natural dynamics, described by Kern[2] as a protein’s “dynamic personality”, can be critical to a protein’s function, and can directly affect the geometry of, for example, an enzyme active site.

A judicious combination of simplified methodologies – rigidity analysis, elastic network modelling, and geometric simulation of flexible motion – can work together to extract valuable information on the large-amplitude, low-frequency motions of a protein structure, at a fraction of the computational cost (in resources and time) of molecular dynamics (MD) approaches[3]. These simplified methods are strongly complementary to both computational MD investigations and experimental biophysical/biochemical studies of protein behaviour.

Recent studies has established that the motions explored by geometric simulation correspond well to the essential dynamics extracted from MD trajectories[4]; that global flexible motions can couple to enzyme active site geometry[5-8]; and, crucially, that a structure generated from geometric simulation of a large amplitude motion can be used as input for further MD investigation[9], as the geometric simulation retains the local bonding geometry and constraint network of the input crystal structure and forbids major bonding distortions and steric clashes.

The information obtained here regarding the flexibility of the enzyme should be of use and interest in structure-based drug design and computational screening of drugs and drug-like molecules capable of binding to and inhibiting the action of the viral protease, such as the pioneering work of Walsh *et al*. reported online within the past few days[10]. The simulations reported here suggest that the protease structure is capable of substantial flexible motions which alter domain orientations, open and close clefts, and affect the geometry of an inhibitor-binding site. The entirety of the computational work described here was carried out in less than one working day on a laptop computer. The inputs and outputs of the simulations, principally consisting of PDB files representing flexible variations on the protease crystal structures, are available for download from the University of Bath research data archive at https://researchdata.bath.ac.uk/772/ [11] with DOI https://doi.org/10.15125/BATH-00772

## Methods, software and data

Input data: crystal structures of a viral protease were downloaded from PDB entries 6Y2E (free protease) and 6LU7 (inhibitor bound).

Visualisations: visualisation and some minor structural editing, *viz*. generation of symmetry copies, removal of water and other heterogroups, removal of alternate sidechain conformations, and renumbering of entries after addition of hydrogens, were carried out with the PyMOL viewer, version 0.99[12]. All structures were aligned to the 6Y2E crystal structure for consistency of view.

Addition of hydrogens, sidechain flipping: the MolProbity[13] web server hosted at Duke University provides online tools carrying out these functions.

Rigidity analysis: carried out using the pebble-game constraint counting algorithm implemented in the software FIRST[14]. FIRST should be available from Arizona State University via flexweb.asu.edu; academic readers should contact the present author if there is difficulty in obtaining the software, as the site appears currently to be down. Recent research has shown that the assignment of energies to polar interactions in FIRST suffers from a bug that leads to some salt-bridge interactions with short donor-acceptor distances being treated as weak rather than strong. Therefore, interactions were identified and energies assigned using a recently developed corrected function, SBFIRST[15]. A copy of the SBFIRST software for constraint identification is included with the deposited dataset for this study.

Normal mode analysis: carried out using the Elnemo elastic network modelling software[16]. This method builds an elastic network model of the protein with one site per residue (using the alpha carbon _CA_ atom coordinates as the node positions), placing springs of uniform strength between each pair of residues lying less than 10 Å apart, and diagonalising the resulting Hessian matrix to identify eigenvectors.

Geometric simulations of flexible motion: carried out using the FRODA geometric simulation engine[17] incorporated into FIRST. The FRODA engine iteratively projects an all-atom model of the protein structure along a normal mode direction[3] while constraining the local geometry to match that of the original protein structure. Frames along each such trajectory are written out as numbered PDB files.

Workflow and protocol: for each protein structure, the complete workflow for this study is as follows.

Download structure in PDB or mmCIF format. In this case, the downloaded structure contains coordinates for a single protein chain, and symmetry operations to generate copies making up the crystal structure.

Visualise structure in PyMOL. Generate symmetry mates. Select the appropriate symmetry mate to make up the homodimer indicated in the PDB entry (under Global Stoichiometry) as the biological entity of interest. Delete other symmetry copies. Alter chain ID(s) in the symmetry copy to give each chain a unique chain ID.

Process structure through the MolProbity website, adding hydrogens at electron-cloud positions, accepting all recommendations for side chain flips. Download the hydrogenated structure (in which all hydrogens have a serial number of 0).

Visualise structure in PyMOL for final cleaning and preparation. Remove alternate conformations (i.e. those with an *altloc* entry which is not a blank or an A). Remove water molecules and extraneous heterogroups such as glycerol, acetyl etc. Save the cleaned structure; in this step, PyMOL renumbers the atoms sequentially, so hydrogens now have appropriate serial numbers.

Identify covalent and noncovalent interactions from the atomic geometry of the protein structure using SBFIRST. This generates lists of covalent bonds, tagged as being either rotatable or non-rotatable (such as the C-N bond in the peptide main chain); hydrophobic tether interactions, wherever two nonpolar sidechains are closely adjacent; and polar interactions, including salt bridges and hydrogen bonds. Polar interactions are assigned energies between 0 and -10 kcal/mol based on the donor-hydrogen-acceptor geometry.

Analyse rigidity with FIRST as a function of hydrogen-bond energy cutoff. This analysis groups atoms into rigid clusters by matching degrees of freedom against constraints. FIRST can carry out a dilution, gradually lowering the cutoff to exclude polar interactions one by one from weakest to strongest. It is also possible to simply carry out “snapshots” of the rigidity at a series of cutoff values, which in this study were -0.0, -0.5, -1.0 … -3.5, -4.0 kcal/mol.

Review constraints based on rigidity information. At this stage, a small number of noncovalent constraints (one in 6Y2E, three in 6LU7) were identifiable as artefacts of crystal packing, and were excluded from the network when carrying out the geometric simulations. Based on the rigidity analysis, a cutoff of -3.0 kcal/mol appeared appropriate for the geometric simulations, in line with previous studies [4, 5].

Normal mode analysis. Extract only the alpha carbon positions and run Elnemo *pdbmat* (generation of matrix) and *diagstd* (diagonalise matrix) functions. The result is a series of normal modes of motion, reported as eigenvector/eigenvalue pairs. When these are sorted by eigenvalue from lowest to highest, the first six modes are trivial rigid-body motions with near-zero eigenvalues (frequencies), while the low-frequency modes from 7 upwards represent flexible motions of the protein itself. This study examines modes 7 to 16, that is, the ten lowest-frequency nontrivial modes, with modes 7 to 12 being shown in the figures.

Geometric simulation of flexible motion. For each normal mode of interest (7 to 16), run FIRST with a chosen energy cutoff (−3.0 kcal/mol) and the previously identified lists of covalent and noncovalent bonding constraints, providing the normal mode eigenvector and a direction of motion (parallel or antiparallel to the vector) along with a directed step size (in this case 0.01Å), and invoking the FRODA engine implemented in FIRST. FRODA carries out a series of steps. In each step, all atom positions are moved along the normal mode direction. Then, the local atomic geometry defined by the bonding constraints and steric exclusion is restored by a few cycles of iterative relaxation. After a user-defined number of steps (in this case 100), a numbered “frame” is written out as a PDB file. A series of such frames describes the trajectory of flexible motion. FRODA continues each run until a user-defined maximum number of steps (in this case 1500) or until “jamming” occurs where the constraints can no longer be satisfied, typically when the motion has led to severe steric clashes through the collision of two domains.

Output data: the inputs, scripts and outputs of the rigidity analysis, normal mode analysis, and geometric simulations for each protein have been deposited in the University of Bath’s data repository and are accessible from https://researchdata.bath.ac.uk/772/ [11]

## Results and discussion

The 6Y2E and 6LU7 crystal structures each contain explicit coordinates for one chain (A) of the protease, and in the case of 6LU7, a chain (C) representing a bound inhibitor. The input structures for analysis and simulation consist of chain A and a symmetry copy (B), making up the biological homodimer. Chain C is likewise duplicated to chain D. The X-ray structure does not contain explicit hydrogens, which are added using MolProbity[13]. With hydrogens added, it is then possible to detect the covalent bonds, polar noncovalent interactions (hydrogen bonds and salt bridges) and hydrophobic-tether interactions using SBFIRST[15] (see Methods). The polar interactions are rated with an effective energy between 0 and -10 kcal/mol based on the donor-hydrogen-acceptor geometry.

Pebble-game rigidity analysis[14, 18] divides the structure into rigid clusters and flexible regions based on the distribution of degrees of freedom and constraints. This rigid cluster decomposition (RCD) depends on the set of polar interactions that are included, based on an energy cutoff that excludes weaker interactions. Information on the relative stability of different portions of the structure is typically obtained at cutoffs around a “room temperature” value of -1 kcal/mol. RCDs are shown in Figure 1. With the inclusion of weaker polar constraints, a single large rigid cluster extends across both chains of the protease dimer. As weaker constraints are eliminated, the large cluster extends across the N-terminal domains (roughly residues 1-200) of both chains, which consist largely of beta-sheet structure, while the largely alpha-helical C-terminal domain breaks up into several smaller rigid clusters, each one a single helix. With further elimination of constraints, the beta-sheet regions become fully flexible.

**FIGURE 1:**
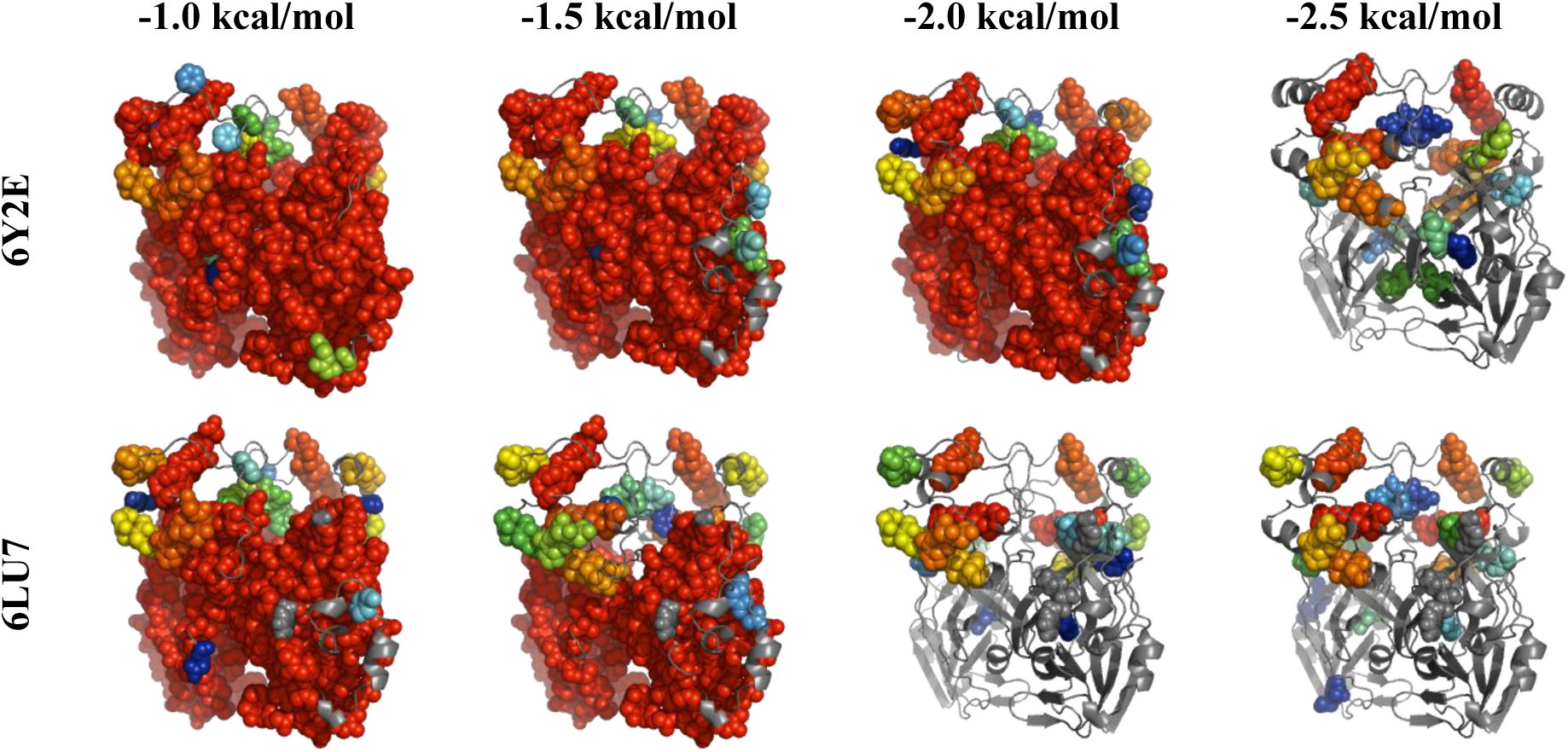
Rigid Cluster Decompositions as a function of hydrogen bond energy cutoff for the empty 6Y2E and inhibitor-bound 6LU7 structures. The twenty largest rigid clusters in each case are shown as spheres and colour-coded from red (largest) to blue; the flexible regions are shown as cartoon.

Flexible motion of a protein structure is best carried out at with a rigidity cutoff such that the structure is fully flexible. The mobility studies that follow are carried out with a cutoff of -3.0 kcal/mol, similar to previous studies on enzymes[4, 5]. It is important to appreciate that even at this lower cutoff the structure is still thoroughly constrained by noncovalent interactions, as illustrated in Figure 2. The core of each folded domain is rich in hydrophobic interactions, as is the interface region between the two beta-sheet domains as the two chains form the homodimer, and arrays of strong polar interactions constrain secondary structure (for example, the backbone hydrogen bonds within alpha helices and beta sheets).

**FIGURE 2:**
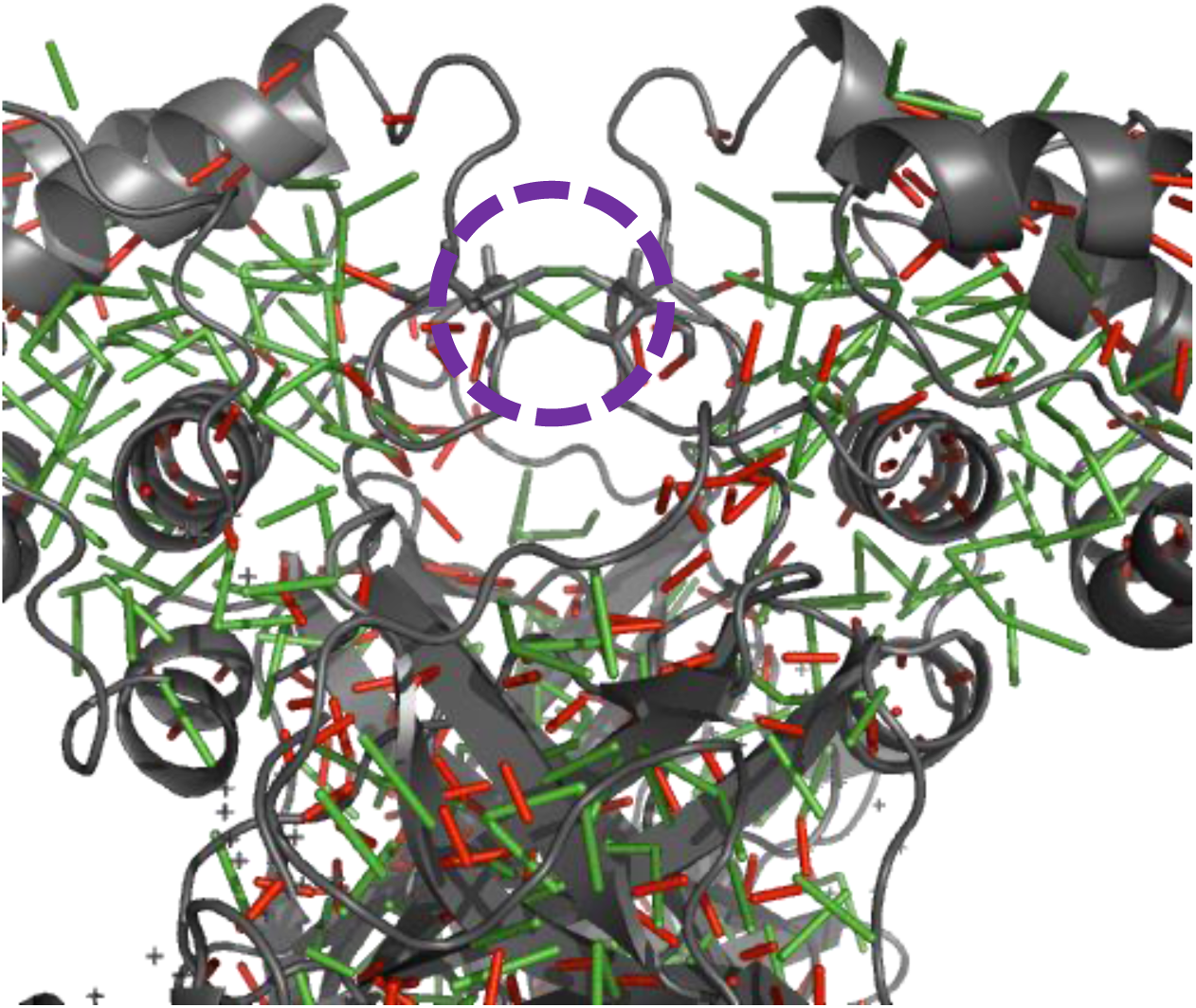
Noncovalent constraints in the 6LU7 structure, shown as green (hydrophobic) and red (polar) lines. The protein mainchain is shown in cartoon representation. The circled area highlights a small set of interdomain hydrophobic constraints, one between the ALA 285 residues of the two chains, and one each between residue THR 280 of one chain and LEU 286 of the other. In the 6Y2E structure only the ALA 285 interaction is present. These interactions may be considered an artefact of crystal packing.

Also highlighted in Figure 2 are a small set of hydrophobic interactions assigned between the N-terminal domains of chains A and B. In the 6Y2E structure, a single tether connects the sidechains of the ALA 285 residues of both chains. In the 6LU7 structure illustrated, there are two additional tethers nearby, each connecting the sidechains of the THR 280 residue of one chain and the LEU 286 of the other. Since the two N-terminal domains are otherwise not connected by noncovalent interactions, and these tethers are not involved in forming a rigid cluster, it seems reasonable to consider these tethers an artefact of crystal packing, and eliminate them from further consideration. Their deletion does not affect the rigid cluster decompositions of the structures. If these tethers were retained, they would somewhat limit (but not entirely prevent) the central cleft-opening motion discussed below.

Normal modes, representing directions of motion intrinsic to the structure, are easily obtained from an elastic network model[16] (see Methods) in which the protein is represented as one site per residue and springs of uniform strength are placed between every pair of sites with a separation less than 10 Å. In the 6LU7 structure, the amino acid residues of the bound inhibitor are included in the elastic network. Of the 3N normal modes of a structure with N residues, six modes of near-zero frequency represent the trivial rigid body motions of the structure, while the low-frequency modes from 7 upwards represent modes of flexible motion. When the structure moves along a low-frequency mode direction, the global geometry changes with minimal change in the local geometry, making these directions of “easy” motion for the protein. The sign of a normal mode eigenvector is arbitrary; motions parallel and antiparallel to each bias direction must be investigated equally.

Linear projection of a protein structure along a normal mode direction would rapidly introduce unphysical distortions into the structure. The geometric simulation approach implemented in FRODA[3, 17] instead applies a bias, moving the structure along a normal mode direction, while maintaining steric exclusion and the bonding geometry and noncovalent constraints of the input structure. This approach rapidly generates physically realistic flexible variations on the input structure which retain all-atom steric and bonding detail.

Geometric simulations of the lowest-frequency nontrivial modes of motion (7 upwards) show that the protease structure is capable of substantial amplitudes of easy motion covering several Ångstroms distance in global root-mean-square displacement (RMSD). In particular, the orientations of the alpha-helical domains relative to the beta-sheet domains can change through flexing and rotation about the interdomain “hinges”, and the beta-sheet domains themselves are large enough to display bending and twisting motions within themselves. These variations are illustrated for the 6Y2E structure in Figure 3 and for 6LU7 in Figure 4.

**FIGURE 3:**
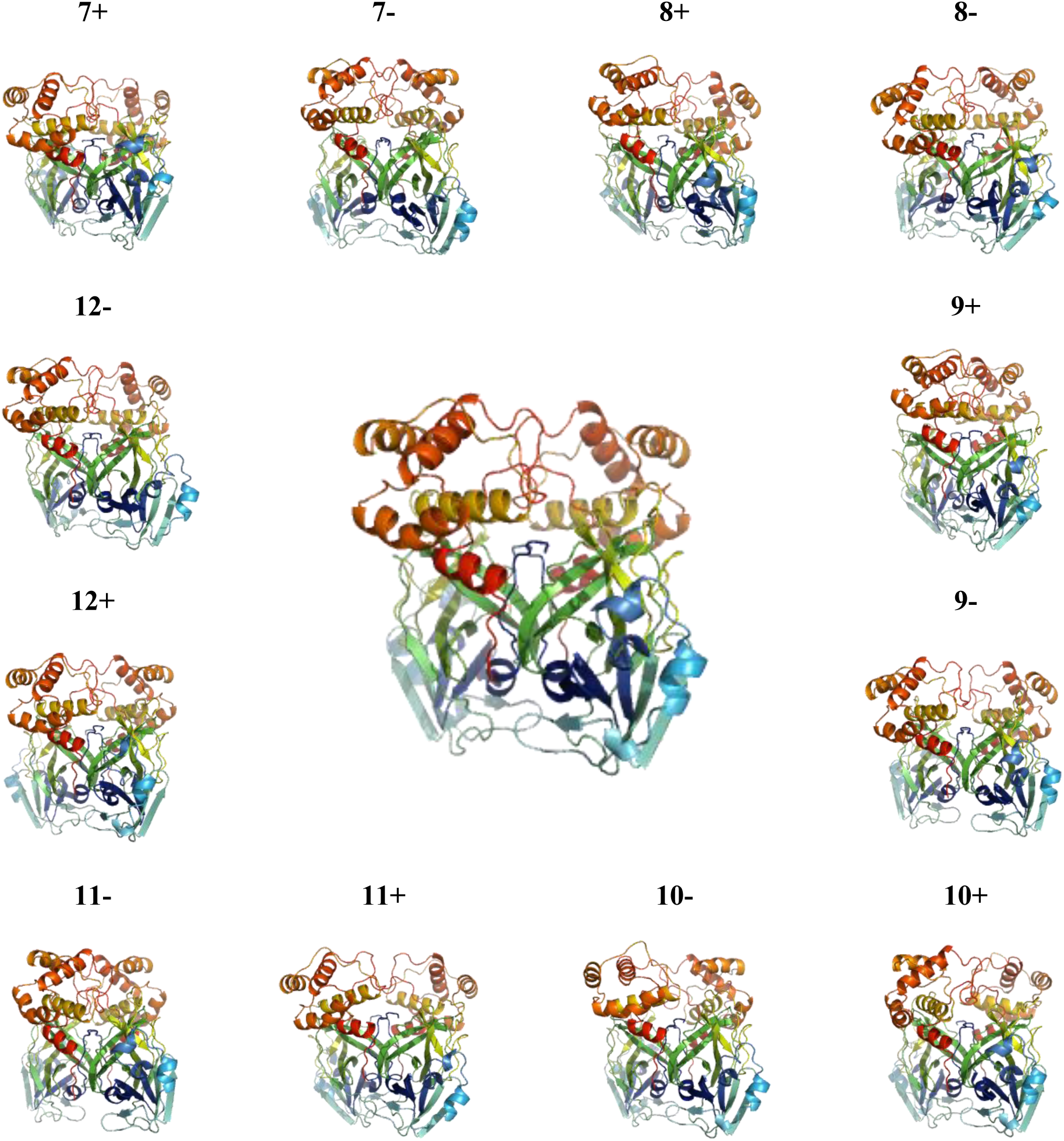
Flexible variations (outer circle) of the 6Y2E structure (centre), shown in cartoon representation and coloured from blue to red along each chain. Each flexible variation is the 1000^th^ frame of a geometric simulation, lying at an all-atom RMSD of 3 to 4 Ångstroms from the starting crystal structure. Note the substantial variations of domain orientation achieved. Each variation is labelled with the normal mode used to bias the geometric simulation, from 7 to 12, and with the direction of bias, parallel (+) or antiparallel (-) to the mode eigenvector, whose sign is arbitrary.

**FIGURE 4:**
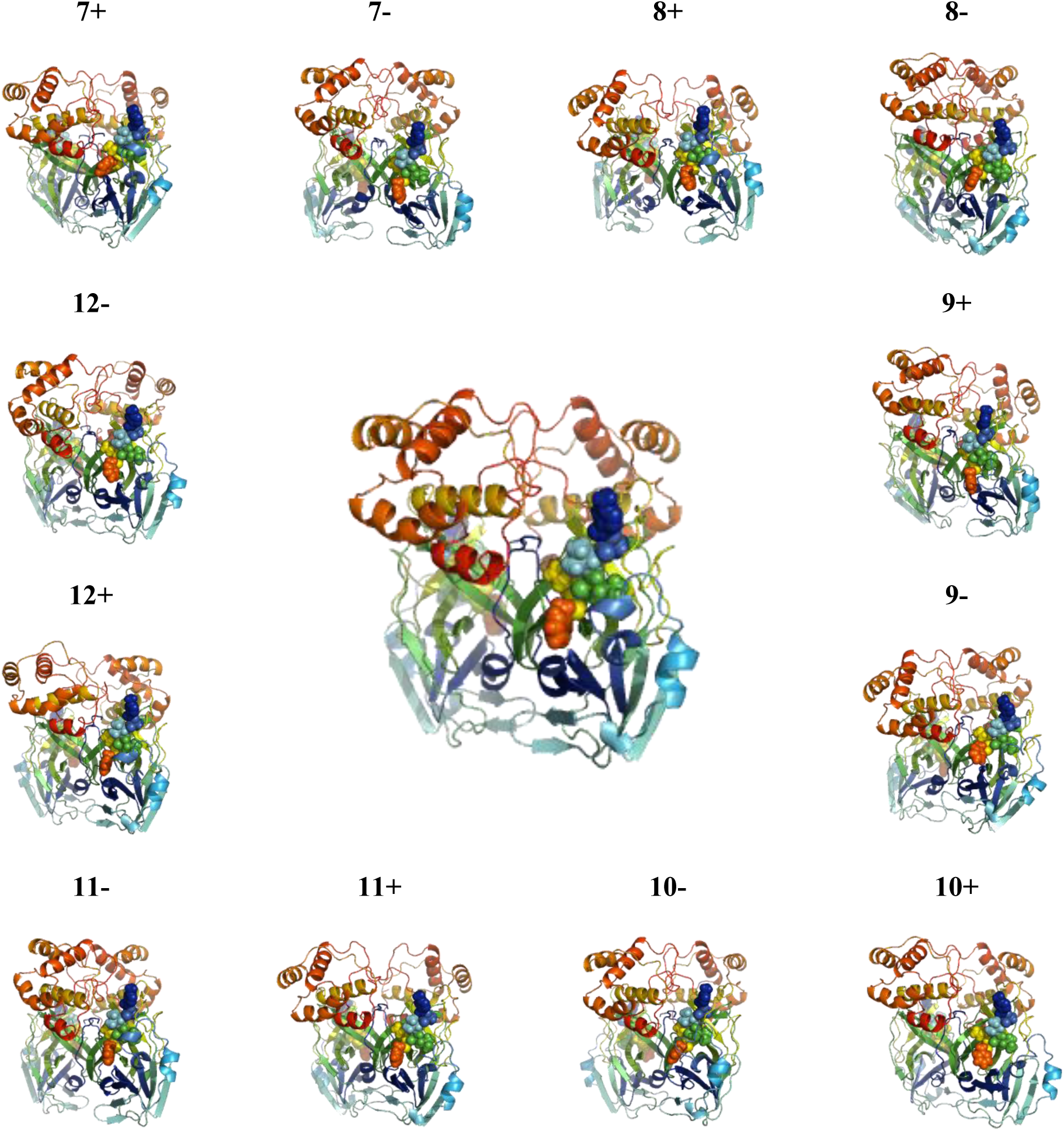
Flexible variations (outer circle) of the 6LU7 structure (centre), shown in cartoon representation and coloured from blue to red along each chain. The bound inhibitor is shown as spheres. Each flexible variation is the 1000^th^ frame of a geometric simulation, lying at an all-atom RMSD of 3 to 4 Ångstroms from the starting crystal structure. Note the substantial variations of domain orientation achieved. Each variation is labelled with the normal mode used to bias the geometric simulation, from 7 to 12, and with the direction of bias, parallel (+) or antiparallel (-) to the mode eigenvector, whose sign is arbitrary.

Flexible variations on a protein crystal structure are potentially relevant to structure-based drug design in at least two ways. Firstly, flexible motion can open or expose clefts and potential binding sites not directly visible in the static crystal structure. Since the protein *in vivo* will be exploring its flexible motion thanks to the Brownian-motion driving force of its solvent (e.g. cytosol), such latent sites may constitute valid target areas for inhibitors.

Secondly, the global low-frequency motion couples to variations in the binding site/active site geometry[5, 6]. Knowledge of the range of flexible variation here is potentially useful for structure-based drug design and/or fragment screening, since attention can be focussed on candidate molecules that interact robustly with the binding site and tolerate its flexibility, in preference to molecules that interact well only with the crystal structure and not with its flexible variations.

Figure 5 illustrates a case of cleft opening. In both the 6Y2E and 6LU7 structures, some of the modes of motion (9- and 11+ in 6Y2E; 8+ and 11+ in 6LU7) have the two alpha-helical domains moving apart from each other, exposing a cleft at the centre of the protein that is a largely covered, narrow tunnel in the crystal structures. If the hydrophobic tethers eliminated previously were retained, the amplitude of this opening motion would be limited, but the cleft opening could still occur in the form of a widening of the tunnel. The protein structures are in this case shown in a space-filling (sphere) representation, since it is the exposure of new surface that is of interest.

**FIGURE 5:**
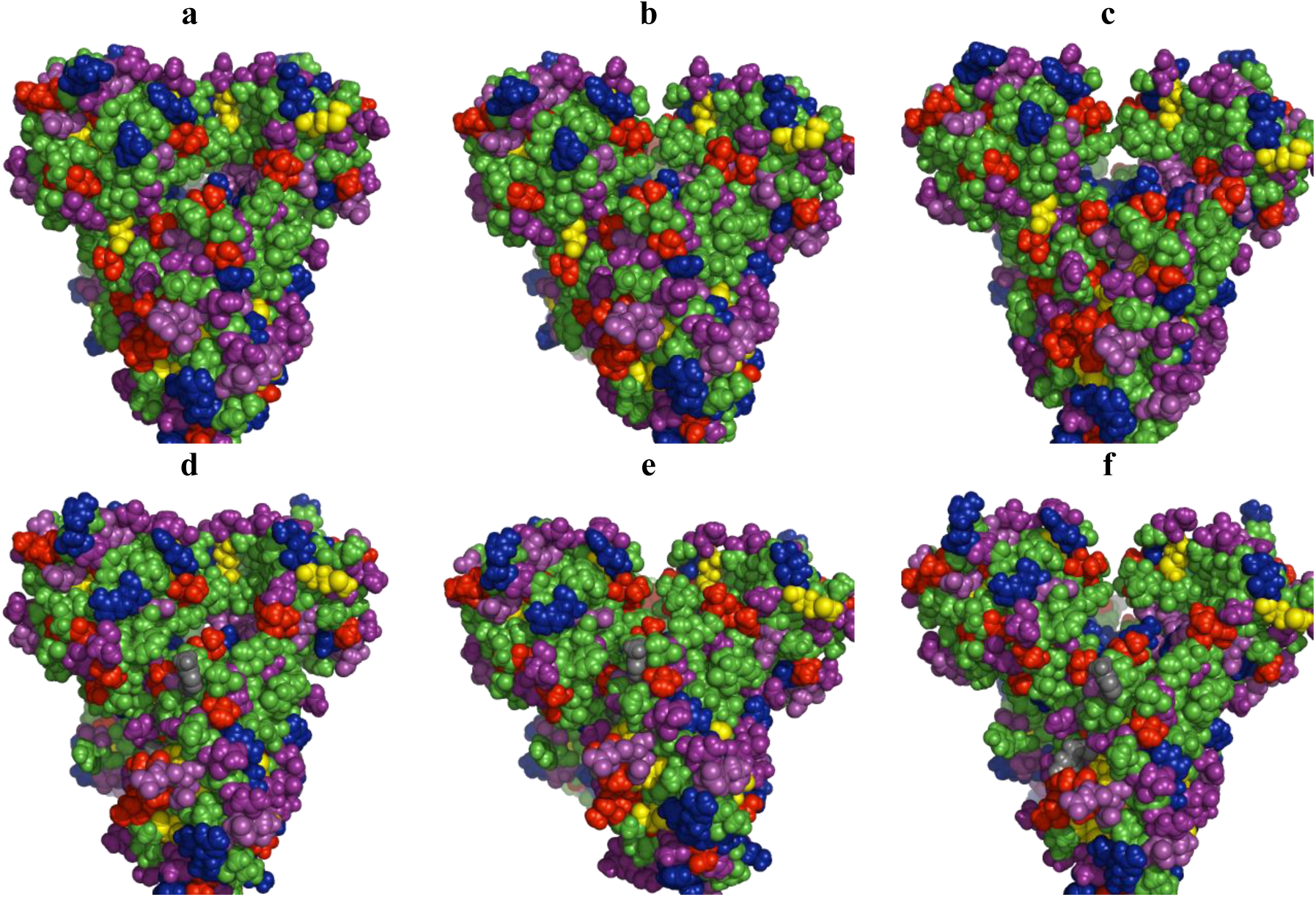
Opening of a cleft between the N-terminal domains during flexible motion. Top row: 6Y2E crystal structure (a) and cleft opening motion along modes 9- (b) and 11+(c). Bottom row: 6LU7 crystal structure (d) and cleft opening motion along modes 8+ (e) and 11+ (f). Protein structure is shown as spheres and coloured by amino acid type. Green, aliphatic or aromatic residues; yellow, cysteine or methionine; red, hydroxyl (serine and threonine); magenta, acidic; purple, amidic; blue, basic.

A more detailed view of this opening cleft is shown in Figure 6. The residues are coloured to bring out the chemical character of the amino acid side chains. The portions of the alpha-helical domain surfaces that move aside to open the cleft display largely hydrophobic residues (aliphatic or aromatic side chains). Intriguingly, the exposed surface area within the cleft is richly lined with a series of basic residues (lysine and arginine), and is flanked by acidic (aspartic and glutamic acids) and polar (threonine) residues. In the crystal structures the basic and acidic residues appear to be involved in a network of inter- and intra-domain salt bridge interactions which stabilise the core of the homodimer. An antagonist capable of targeting this zone of strongly polar surface geometry exposed by flexible motion could potentially disrupt the dynamics of the enzyme and interfere with its function.

**FIGURE 6:**
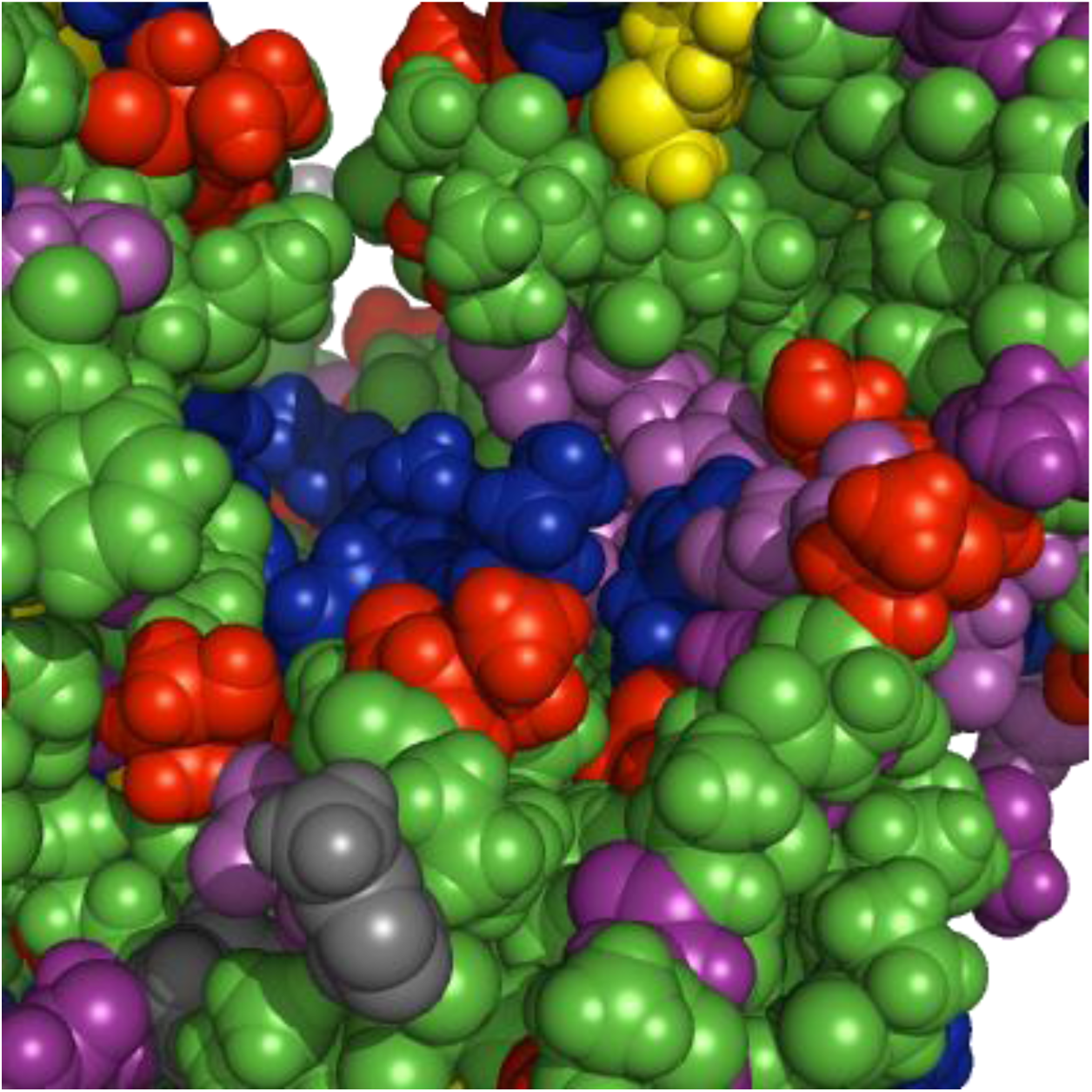
Detail view of interdomain cleft opening (structure 6LU7, mode 11+, 1000^th^ frame of geometric simulation). Protein structure is shown as spheres and coloured by amino acid type. Green, aliphatic or aromatic residues; yellow, cysteine or methionine; red, hydroxyl (serine and threonine); magenta, acidic; purple, amidic; blue, basic. One end of the bound inhibitor is visible as grey spheres in the lower left. The exposed cleft is rich in basic, acidic and polar residues.

In the 6LU7 structure, a binding site on the flank of each beta-sheet domain is occupied by the inhibitor N-((5-methylisoxazol-3-yl)carbonyl)alanyl-L-valyl-N∼1∼-((1R,2Z)-4-(benzyloxy)-4-oxo-1-(((3R)-2-oxopyrrolidin-3-yl)methyl)but-2-enyl)-L-leucinamide (chain C of the structure), hereinafter “N3”. This inhibitor is listed in the PDB as PRD_002214; it is a peptide-like inhibitor, with a central core of amino acids modified at either end with heterogroups, and was previously known to inhibit a protease of a feline coronavirus[19] (see PDB entry 5EU8). In the 6Y2E structure the same binding cleft is empty, which means that its geometry is likely to change in the course of flexible motion of the beta-sheet domain.

Figure 7 shows the N3 binding site of the 6LU7 structure and the corresponding region of the 6Y2E structure. As noted, all structures have been aligned to the 6Y2E crystal structure for ease of viewing and comparison. Figure 7 also shows flexible variations of the 6Y2E site in the course of flexible motion. A set of residues forming a rough square around the binding site (GLU 166, GLN 189, ASN 142 and THR 25) are highlighted in white as a guide to the eye. It is immediately visible that openings, closings and flexible distortions of the site are an intrinsic feature of the flexible motion of the protease. Note in particular the effect of motion along the lowest-frequency nontrivial mode direction, mode 7. Motion biased antiparallel to the mode eigenvector (7-) includes an opening of the cleft, while motion biased parallel to it (7+) closes it. Since the N3 inhibitor is quite deeply embedded in the binding site of the 6LU7 structure, opening and closing of this cleft in the course of the protein’s natural motion may be necessary for molecules to access the cleft. On visual inspection, the site in the 6Y2E structure is clearly slightly more closed that in 6LU7, consistent with the proposition that such opening/closing motions are intrinsic to the protein.

**FIGURE 7:**
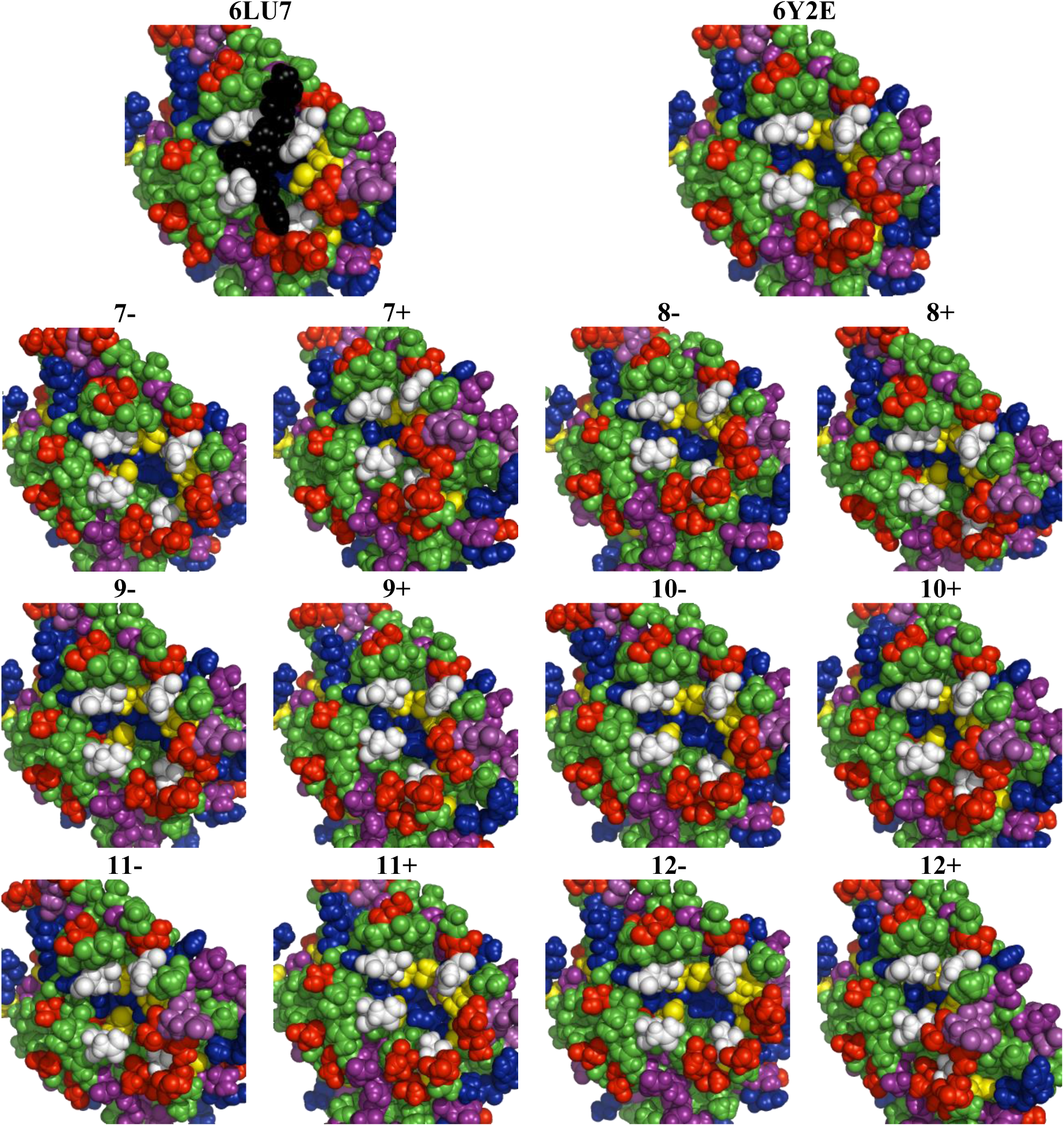
Top row: the N3 inhibitor binding site of the 6LU7 structure, and the corresponding empty site of the 6Y2E structure. Rows below: flexible variations of the 6Y2E site from geometric simulations along normal modes as indicated. Residues 1-200 of a single chain of the protease are shown as spheres and coloured by amino acid type. Green, aliphatic or aromatic residues; yellow, cysteine or methionine; red, hydroxyl (serine and threonine); magenta, acidic; purple, amidic; blue, basic. To highlight the binding site, residues GLU 166, GLN 189, ASN 142 and THR 25 are coloured white. Substantial opening and closing of the binding site cleft occurs during the flexible motion.

## Conclusions

The rigidity and flexibility of two recently reported crystal structures (PDB entries 6Y2E and 6LU7) of a protease from the SARS-CoV-2 virus, the infectious agent of the COVID-19 respiratory disease, has been investigated using pebble-game rigidity analysis, elastic network model normal mode analysis, and all-atom geometric simulations. This computational investigation of the viral protease follows protocols that have been effective in studying other homodimeric enzymes. The protease is predicted to display flexible motions *in vivo* which directly affect the geometry of a known inhibitor binding site, *e*.*g*. through an opening/closing motion, and which open new potential binding sites elsewhere in the structure. A database of generated PDB files representing natural flexible variations on the crystal structures has been produced and made available for download from an institutional data archive. This information may inform structure-based drug design and fragment screening efforts aimed at identifying specific antiviral therapies for the treatment of COVID-19.

## Acknowledgements

SAW has received funding from the European Research Council (ERC) under the European Union’s Horizon 2020 research and innovation programme (grant agreement No 648283 “GROWMOF”, PI: Prof. T. Düren).

## References

1. WHO, Naming the coronavirus disease (COVID-19) and the virus that causes it. 2020.

2. Henzler-Wildman, K. and D. Kern, Dynamic personalities of proteins. Nature, 2007. 450(7172): p. 964–72.

3. Jimenez-Roldan, J.E., et al., Rapid simulation of protein motion: merging flexibility, rigidity and normal mode analyses. Physical Biology, 2012. 9(1).

4. Wells, S.A., et al., Structure and Function in Homodimeric Enzymes: Simulations of Cooperative and Independent Functional Motions. Plos One, 2015. 10(8).

5. Jones, H.B.L., et al., A complete thermodynamic analysis of enzyme turnover links the free energy landscape to enzyme catalysis. Febs Journal, 2017. 284(17): p. 2829–2842.

6. Jones, H.B.L., et al., Exposing the Interplay Between Enzyme Turnover, Protein Dynamics, and the Membrane Environment in Monoamine Oxidase B. Biochemistry, 2019. 58(18): p. 2362–2372.

7. Li, H.L., et al., Protein flexibility is key to cisplatin crosslinking in calmodulin. Protein Science, 2012. 21(9): p. 1269–1279.

8. Adjogatse, E., et al., Structure and function of l-threonine-3-dehydrogenase from the parasitic protozoan Trypanosoma brucei revealed by X-ray crystallography and geometric simulations. Acta Crystallographica Section D-Structural Biology, 2018. 74: p. 861–876.

9. Romer, R.A., et al., The flexibility and dynamics of protein disulfide isomerase. Proteins-Structure Function and Bioinformatics, 2016. 84(12): p. 1776–1785.

10. Walsh, M., Main protease structure and XChem fragment screen. 2020.

11. Wells, S.A., Dataset for “Rigidity, normal modes and flexible motion of a SARS-CoV-2 (COVID19) protease structure”. 2020.

12. LLC, S., The Pymol molecular graphics system. 2010.

13. Williams, C.J., et al., MolProbity: More and better reference data for improved all-atom structure validation. Protein Science, 2018. 27(1): p. 293–315.

14. Jacobs, D.J., et al., Protein flexibility predictions using graph theory. Proteins-Structure Function and Genetics, 2001. 44(2): p. 150–165.

15. McManus, T.J., S.A. Wells, and A.B. Walker, Salt bridge impact on global rigidity and thermostability in thermophilic citrate synthase. Physical Biology, 2020. 17(1).

16. Suhre, K. and Y.H. Sanejouand, ElNemo: a normal mode web server for protein movement analysis and the generation of templates for molecular replacement. Nucleic Acids Research, 2004. 32: p. W610–W614.

17. Wells, S.A., et al., Constrained geometric simulation of diffusive motion in proteins. Physical Biology, 2005. 2(4): p. S127–S136.

18. Wells, S.A., J.E. Jimenez-Roldan, and R.A. Romer, Comparative analysis of rigidity across protein families. Physical Biology, 2009. 6(4).

19. Wang, F., et al., Crystal Structure of Feline Infectious Peritonitis Virus Main Protease in Complex with Synergetic Dual Inhibitors. J Virol, 2016. 90(4): p. 1910–7.

